# SLTRiP induces long lasting and protective T-cell memory response

**DOI:** 10.1101/2021.01.07.425694

**Authors:** Afshana Quadiri, Mohammad Kashif, Inderjeet Kalia, Agam Prasad Singh

**Author notes:** To whom correspondence should be addressed: Agam P. Singh, National Institute of Immunology, Aruna Asaf Ali Marg, New Delhi-110067, India. & 14 26715032, (APS).

## Abstract

Major developments have been made in the past many years to characterize and explore potential vaccine candidates that can induce host immune responses against parasite. These advances were based on the fact that the induction of host immune responses could effectively target parasite at different stages of its life cycle and thus, abrogate *Plasmodium* infections. The role of T-cells against malaria comes from initial studies on rodents showing these cells could inhibit parasite development during pre-erythrocytic stages. Since then, the importance of the cellular immune responses against malaria has been increasingly emphasized, especially for vaccine development against pre-erythrocytic stages. Previous work in our laboratory has confirmed that SLTRiP confers protection against the pre-erythrocytic stage of *Plasmodium* growth in rodents. Here we report that the protection is mainly due to cell mediated immune responses and *Pb*SLTRiP specific cellular memory responses could be efficiently recalled in mice challenged with *P. berghei* parasites even after a year following immunization. Our results thereby, highlight the role of the T cell response involved in protection. Characterization of T-cells by intracellular cytokine staining (ICS) revealed that the induced T cells were polyfunctional and involved in secretion of pro-inflammatory cytokines which mediate anti-parasitic activity. The findings contribute to our understanding of the immunological mechanisms underlying the protective vaccines.

## 1. Introduction

Malaria is a deadly mosquito-borne infectious disease affecting millions of people each year. Approximately 228 million cases of malaria occurred in 2018 and an estimated 405,000 related deaths as reported in World Malaria Report 2019 [1]. The disease manifests itself in a broad range of symptoms which include fever, headache, vomiting, and dehydration. Untreated malaria can even lead to severe anemia, multiple organ failure, convulsions, coma, and even death. Although, there has been considerable success in reducing malaria mortality rates in the last 20 years; the burden of disease continues to exist. Due to its extraordinary biological complexity human *Plasmodium spp.* have been developing resistance against all the common anti-malarial drugs [2, 3]. An increasing resistance of mosquitoes against insecticides has also been reported [4, 5]. These concerns feature the requirement of developing an efficient vaccine against malaria.

The vaccination approach using Radiation Attenuated Sporozoites (RAS) showed that it is possible to attain the sterile immunity against malaria [6, 7]. The response was largely directed towards sporozoite and hepatic stages, affecting the subsequent stages also [8, 9]. RAS has been paramount in studying the immune responses generated during pre-erythrocytic parasite invasion [10, 11]. RAS immunizations showed the major role of T cells in protection against sporozoite challenge via adoptive transfer of T-cells and T-cell depletion experiments [7, 12, 13]. Although, both humoral and cell-mediated responses play an important role in controlling the disease, the existing research points at higher potential of T cells in inhibiting the growth and development of pre-erythrocytic *Plasmodium* and thereby reducing the magnitude of the parasite load entering the erythrocytic cycle [14–16] T-cells generate protective responses against many other pathogens which include HIV, influenza virus, CMV and many tumors; suggesting that T cell induction could provide prophylactic efficacy against malaria. T-cells form clusters around infected hepatocytes and secrete cytokines which include IFN-γ, TNF-α, TNF-related apoptosis-inducing ligand (TRAIL), and perforin [17, 18]. These cytokines eliminate infected hepatocytes in multifactorial manner. The literature points at protective immunity mediated by T cells and demonstrates the possible role of stimulated T-cells in response to *Plasmodium* proteins, against liver-stage infection.

A number of vaccine candidates against malaria are undergoing clinical trials and RTS,S has been made commercially available [19–22]. This vaccine is based on parasite coat protein Circumsporozoite Protein (CSP). It contains a part of CSP sequence co-expressed and fused to Hepatitis B surface antigen and formulated along with the chemical adjuvant AS01 to enhance immune response [23]. The results from Phase III clinical trials of RTS,S has revealed that although individuals, especially children, presented high levels of neutralizing antibodies towards CSP after vaccination, it conferred only 40-50% protection [24–26]. Evaluation of RTS,S immune responses, showed it generated high antibody titers and moderate CD4^+^ responses [27]. Although these antibodies inhibit the number of sporozoites infecting hepatocytes, the immune studies using RAS have shown that the pre-erythrocytic protection is primarily dependent on CD8^+^ T cells with contribution from antibodies, CD4^+^ cells, gammadelta T cells and natural killer (NK) cells [16, 28, 29]. RAS mediated protection was abrogated when mice infected with malaria were depleted of CD8^+^T [12, 30]. Hepatic cells express parasitic peptides on their MHC Class I inducing cytotoxic T cells and their associated immunity [31]. In addition, hepatic APCs also express exogenous parasitic antigens on MHC class II and cross present via MHC class I [32]. The property of Kupffers cells and hepatocytes to present antigen to CD8^+^ T-cells is the major reason of protection attributed to attenuated parasite [33, 34].

The T-cells targeting the pre-erythrocytic stage of *Plasmodium* could be induced by irradiated or attenuated sporozoites as well as several subunit vaccines. The current literature on anti-malarial immune responses suggests that T cell responses against multiple antigenic targets may be key for the development of a highly efficacious malaria vaccine [35]. However very little is known about how these protective T-cells are activated during parasite liver stages. In addition, a large number of T-cells are required for protective efficacy [36]. One reason could be that the infection percentage is so small for T-cells to be able to scan and identify all infected hepatocytes. The development of liver stage of parasite was inhibited by CD8^+^ T cells produced against defined epitopes of *Plasmodium berghei* and *Plasmodium yoelii* circumsporozoite (CS) protein[15]. Subsequently, effector CD8^+^ T cells have been induced in transgenic mice expressing a T-cell receptor specific for the MHC l restricted epitope of the CS of *P. yoelii[37].* Heterologous prime-boost immunization regimes have provided a means of inducing and studying T-cell immunity in pre-clinical models.

Our previous work, reported the protective efficacy of a novel antigen SLTRIP [38]. The SLTRiP immunization affected the growth of parasites in hepatocytes through delay in the prepatent period of parasite by 3-4 days. Immunized mice showed high level of protection post sporozoite challenge and exhibited 10,000 fold less parasite 18srRNA copy numbers in liver, emphasizing the vaccine potential of SLTRiP [38]. Furthermore, the subsequent results demonstrated that the protective efficacy by SLTRiP is localized in conserved central region. The peptides designed from such region showed protective efficacy equivalent to whole protein [39]. These data support the potential of the *P. berghei* protein SLTRiP and its orthologues in human malaria parasite, as a target antigen for malaria vaccine development. However this protection was primarily through the cellular arm of the immune system as the anti-SLTRiP antibodies could not neutralize the parasite *in vitro.* In view of our earlier results, the study aims to investigate cellular immunological and memory responses induced by *Pb*-SLTRiP protein.

## 2. MATERIALS AND METHODS

### 2.1) Experimental animals and parasites

The C57BL/6 male/female mice around 6-8 weeks old, were used in all animal experiments. The animal work was conducted in accordance with NII’s Institutional Animal Ethics Committee (IAEC) rules. The IAEC approval number for the project is NII-312/13. The mice were injected with ketamine/xylazine intra-peritoneally for short-term anesthesia. At the end of each experiment the mice were sacrificed humanely by cervical dislocation.

### 2.2) Parasite Cycle

Six to eight week-old male/female C57BL/6, BALB/C and CD1 mice were used for growing parasites. *Plasmodium berghei* ANKA parasites were cycled between mice and *Anopheles stephensi* mosquitoes. Mosquitoes were starved overnight and fed on infected mice. These infected mosquitoes were maintained at 19°C and 70 –80% relative humidity and fed on cotton pads soaked in 20% sucrose solution for 18 days post infected blood meal feeding. After 18 days sporozoites were extracted from dissected salivary glands of infected mosquitoes. For this infected mosquitoes were first rinsed with 50% ethanol, washed twice with 1X PBS, and dissected in RPMI 1640 media containing 10% FBS. To obtain sporozoites, salivary glands were ground gently and centrifuged at 800 rpm for 4 min to remove mosquito tissue. The number of sporozoites present was then determined by counting in a hemocytometer.

### 2.3) Immunization with purified SLTRiP Protein Fragments and SLTRiP peptides

Groups of C57BL/6 mice (SLTRiP and control immunized, 6 mice/group) aged 6 –8 weeks were immunized. Priming was done with 50μg of protein in complete Freund’s adjuvant (Sigma, India) per mouse. In the three subsequent boosters the amount of protein used was 25 μg per mouse mixed with incomplete Freund’s adjuvant (Sigma, India). Boosts were given on days 15 and 28 post-priming. The control group was immunized in an identical manner with GST protein.

### 2.4) Enzyme Linked ImmunoSorbent Assay (ELISA)

Culture supernatants from in vitro stimulated splenocytes were collected after 60 hours of incubation in a CO_2_ incubator. Secreted cytokines were measured by ELISA using a ebiosciences kit following manufacturer’s instructions. The purified anti cytokine antibody was added to the wells of enhanced protein binding ELISA plate, sealed and incubated at 4^0^C overnight. Next day the antibody solution was removed and the plate was blocked using blocking buffer for 1-2 hours at room temperature (RT) to prevent non-specific binding. Plate was washed 3 times with 1X PBST. Biotinylated anti-cytokine detection antibody was added to plate, sealed and incubated at RT for 1 hour. It was washed again 3 times with 1X PBST. Secondary antibody conjugated with HRP was added to the wells, sealed and incubated again at RT for 30 min. Plate was washed 5 times with 1X PBST developed using TMB substrate until color starts to appear. Optical density was measured at 450 nm in a microplate reader.

### 2.5) PBMCs isolation

To isolate PBMCs, heparinized whole blood was collected from mice. The blood was diluted with 1XPBS and gently layered over an equal volume of Ficoll in a Falcon tube. The tube was centrifuged for 30 minutes at 400g without brake. The white and cloudy layer of PBMCs formed was collected. The PBMCs were washed twice with PBS at 300g. The percentage viability was estimated using Trypan blue staining.

### 2.6 Splenocyte isolation

The spleen was removed from mice, washed with PBS and crushed between the frosted ends of a sterile slide. The suspension was passed through a cell strainer. The splenocytes obtained were treated with RBC lysis buffer, washed and cultured in complete media. The percentage viability was estimated using Trypan blue staining before culturing.

### 2.7 Intracellular cytokine staining assay

Splenocytes from immunized and control immunized mice were collected and stimulated with SLTRiP protein(10ug/ml) or PMA/I or medium alone in the 96-well culture plates at 37°C in a humidified chamber with 5% CO2 for 12 h. The secretion inhibitor Brefeldin A was added at a concentration of 10ug/ml, and the plates were incubated further for 12h at 37°C. At 24h, the cells were collected and immunolabeled with surface markers using PerCP-CD3, permeabilised and immunolabeled for intracellular cytokines using FITC anti-IFNγ, and PE-TNFα, monoclonal antibodies. After labeling, the cells were washed, fixed with 1% paraformaldehyde, and kept in the dark at 4°C until acquisition. The data acquisition was carried out on a BD verse using FACS Diva software. One-thousand events were acquired for all tests.

## 3. Results

### 3.1 Immunogenicity of SLTRiP protein in mice

The cellular immune responses are known to be essential for protection against liver stage of *Plasmodium* parasite. To continue with these studies, the cellular responses induced by SLTRiP protein were investigated in detail. In order to determine the generation and magnitude of antigen specific T-cell responses against SLTRiP immunization, C57BL6 mice were immunized with SLTRiP and analyzed, corresponding to peak in immunogenicity. The mice immunization followed the prime-boost regimen (Fig 1). To characterize the T-cell responses, splenocytes and peripheral blood mononuclear cell (PBMC) were isolated from SLTRiP immunized and control immunized mice. The cells were collected, stimulated by SLTRiP *in vitro* and analyzed by FACS for presence antigen specific T-cells. For this, the expression of activation marker CD69 on CD3^+^ T-cells was analyzed (Fig 2A). A higher percentage of CD3^+^CD69^+^ T-cells in SLTRiP immunized mice splenocytes was observed. The SLTRiP specific CD3^+^T-cells were efficiently recalled *in vitro* when splenocytes and PBMCs were stimulated *in vitro* with SLTRiP, leading to their proliferation and an increase in CD69 marker. The results of percentage of activated T-cells obtained from all mice were quantified and corresponded to a significant increase in SLTRiP immunized mice compared to control (Fig 2B and 2C).

**I).**
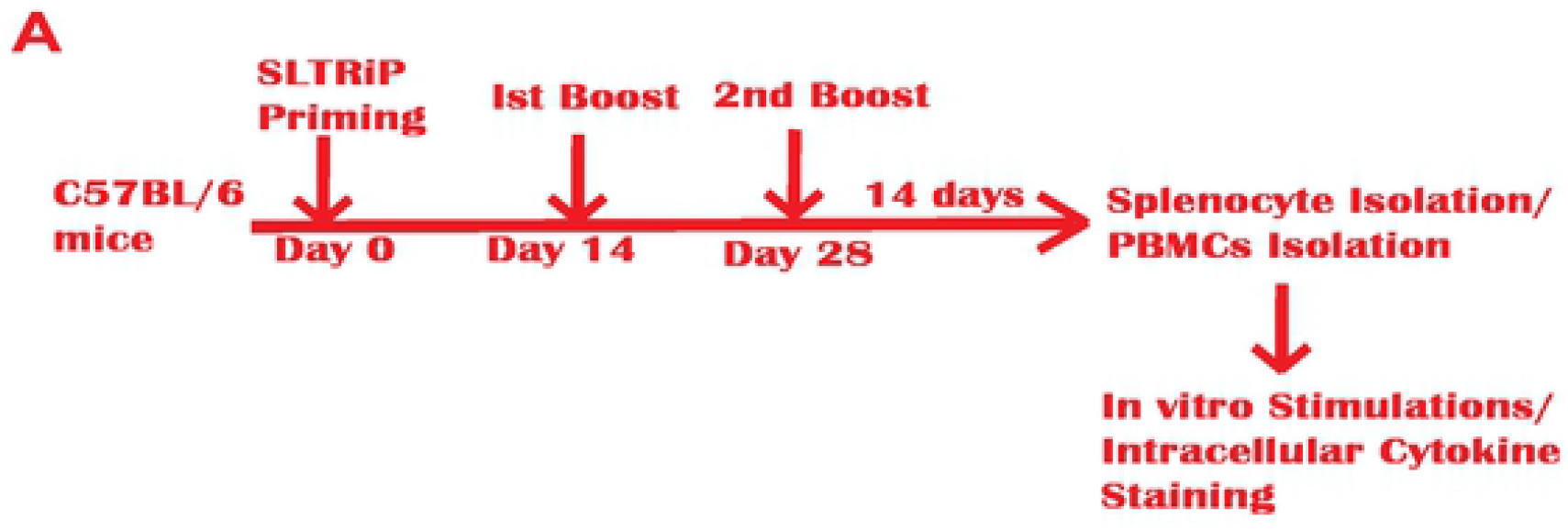
Mice immunization for splenocyte/PBMCs isolation. C57BL6 mice were immunized with SLTRiP/ Control following the prime-boost regimen. Splenocytes and PBMCs were isolated from immunized mice and used for in vitro protein stimulation or Intracellular Cytokine Staining assays.

**II).**
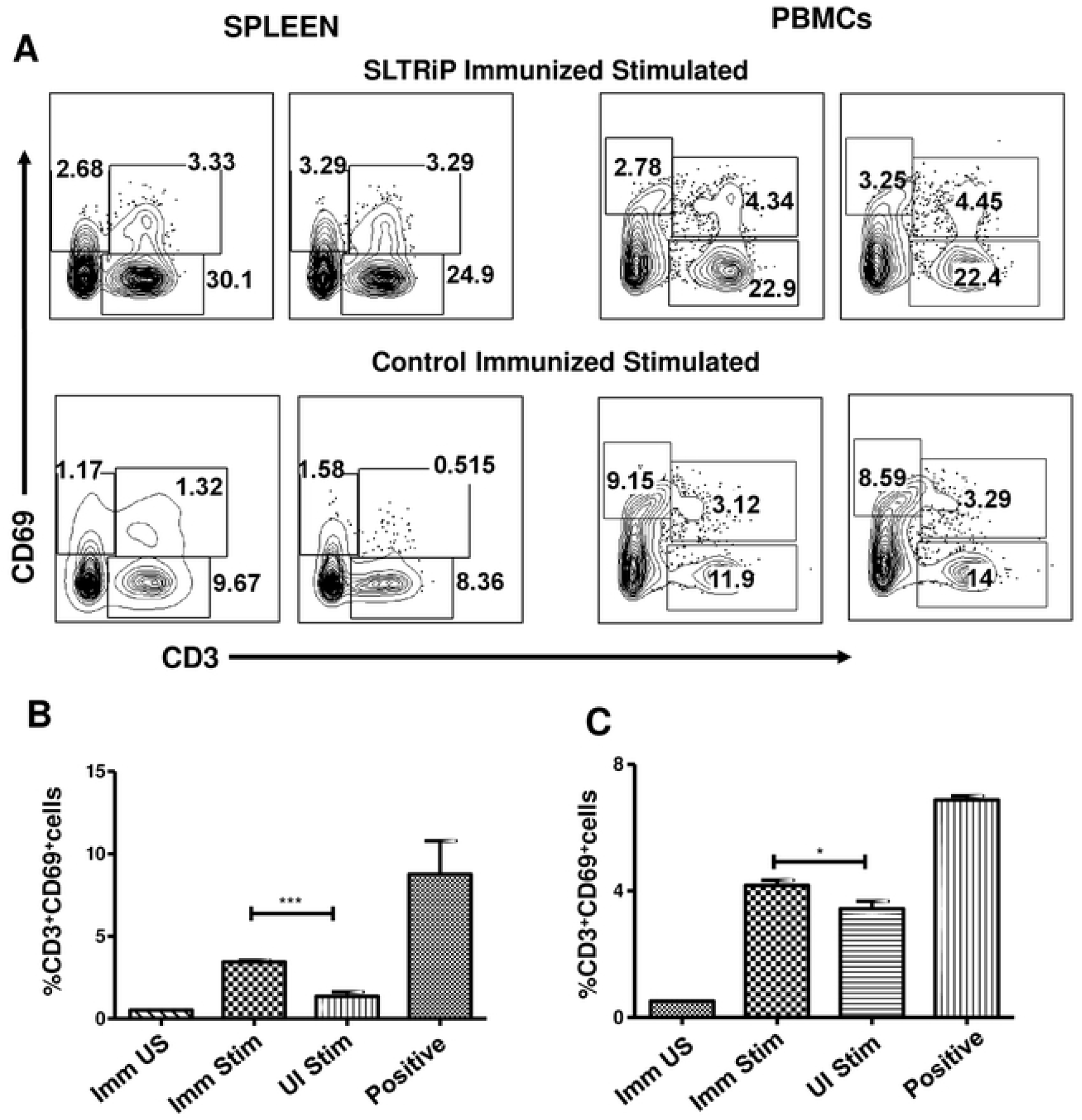
Antigen Specific T-Cell Responses Against SLTRiP Protein. Splenocytes and PBMCs were isolated and stimulated with antigen-SLTRiP, in presence of interleukin (IL)-2. The cells were stained with monoclonal antibodies against cell surface CD3 and CD69 marker and analysed by flow cytometry. The figure represents the total percentage of activated or CD69^+^ cells among a total of CD3^+^ cells for antigen stimulated sample from SLTRiP immunised and control immunised mice (A). The size of the expanded activated T-cell population post *in vitro* stimulation, measured by flow cytometry was plotted. The figure shows the percentage of CD3+CD69+ or dual positive cells in spleen (B) and PBMCs (C). All data are means and standard errors (SEs) based on four mice per group. ***P < 0.0005 by unpaired T-test. ImmUS: Immunised Unstimulated; Imm Stim: Immunised Stimulated; UI Stim: Control Immunised Stimulated; Positive: PMA/I used as positive control.

CD69 is an early lymphocyte activation marker that is expressed on the surface of lymphocytes when the latter encounter antigen presenting cells. The increase in CD3^+^CD69^+^ cells demonstrate that the antigen specific cells have recognized antigen SLTRiP in vitro and were activated in response to it. CD69 marker is increased in responding CD3^+^ T-cells as the latter encounter specific antigen; indicating that SLTRiP protein was able to prime T-cells when mice were immunized with same. These results thus, point at the occurrence or presence of functional T-cell epitopes within the protein, responsible for protection that comes from SLTRiP protein.

### 3.2 Characterization of the polyfunctionality of the induced T cell responses

Pro-inflammatory cytokines like IFN-γ and TNF-α are of key importance against malaria. They are associated with effective host defense against intracellular pathogens. These cytokines allow the accumulation of activated CD8^+^ T-cells and drive their differentiation into functional effectors. In this regard, the cytokine secretion and polyfunctionality of induced T-cell responses were accessed because polyfunctional T-cells are correlated with protective efficacy. For this, splenocytes were taken from SLTRiP immunized and control immunized mice (following prime-boost regimen Fig1), cultured and stimulated with SLTRiP. The supernatant was collected and analyzed for IFN-γ (Fig 3A) and TNF-α (Fig 3B) by ELISA. Cytokine quantitation in stimulated splenocytes, from SLTRiP immunized mice and control, has shown simultaneous induction of both cytokines.

**III).**
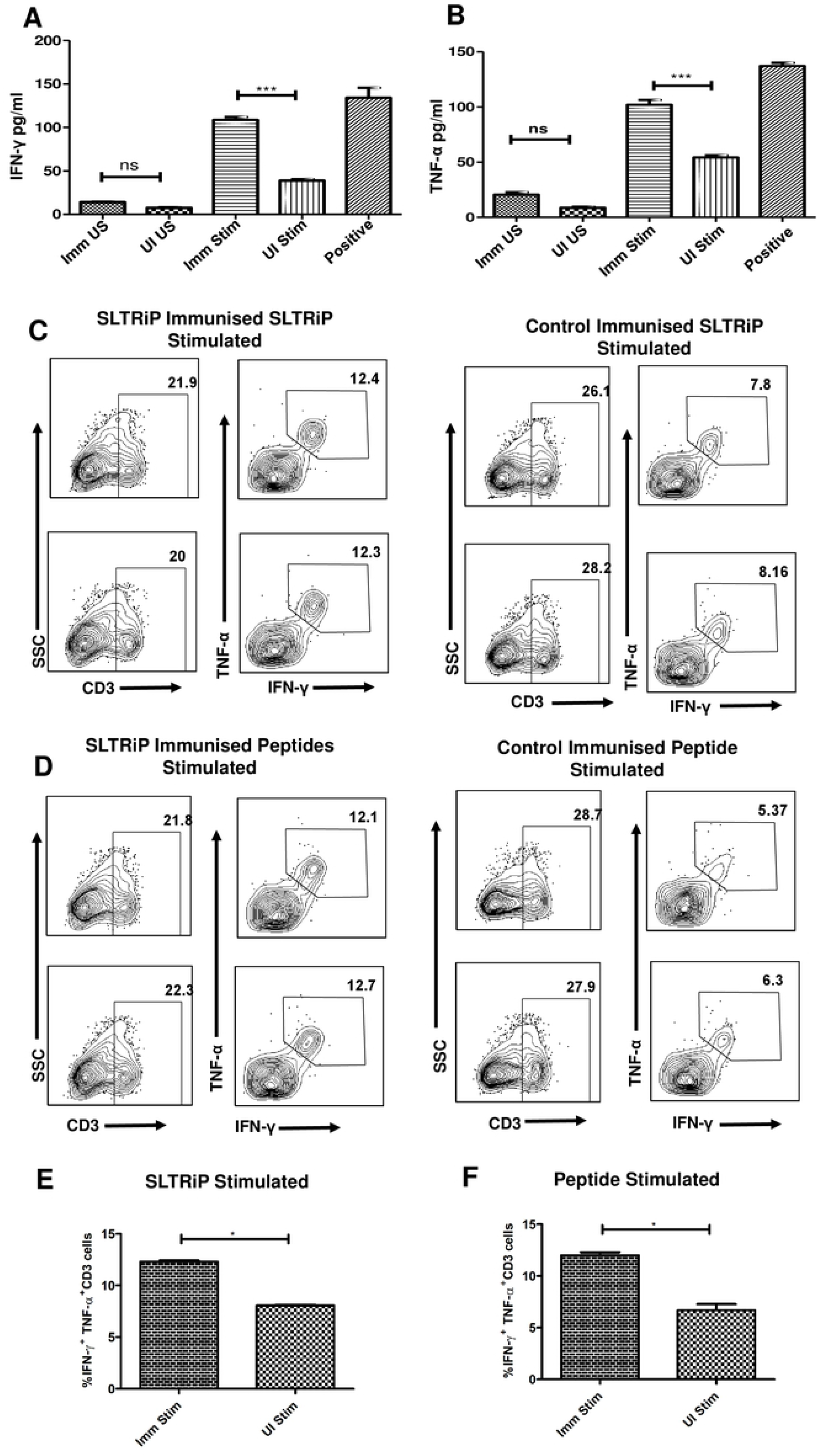
Polyfunctionality of the induced T cell responses: The splenocytes isolated from SLTRiP immunized and control immunized mice were stimulated with antigen-SLTRiP, in presence of interleukin (IL)-2. The cell culture supernatants were collected and the levels of IFN-γ (A) and TNF-α (B) were determined by ELISA. Data have been presented as mean ± SEM. ***P < 0.001; ns, non-significant; by One way ANOVA using Tukeys Multiple comparison test. Imm US: Immunized Unstimulated: UI US: Control Immunised Unstimulated; Imm Stim: Immunised Stimulated; UI Stim: Unimmunised Stimulated; Positive: PMA/I as positive control. Splenocytes from SLTRiP immunized and control were cultured and stimulated with SLTRiP *in vitro* (C) or with T-epitoe peptides (D) in presence of protein transport inhibitor and analyzed by flow cytometry for monofunctional and polyfunctional T-cells (C, D). The increase in the frequency of double positive (IFN-γ and TNF-α) SLTRiP stimulated cells (E) or T-epitoe peptides stimulated (F), has been plotted. Data have been presented as mean ± SEM. *P < 0.05; using Mann Whitney test. Imm Stim: Immunised Stimulated; UI Stim: Control Immunised Stimulated.

The determination of antigen specific cytokine responses of T-lymphocytes after immunization is difficult due to low frequency of responder cells. In order to analyze these responses and observe if these responses were coming from same or different cells we used intracellular cytokine staining assay. The assay was performed using splenocytes from immunized mice. The cells were immunolabelled against surface marker CD3 and against intracellular cytokines IFN-γ, and TNF-α using monoclonal antibodies. Cytokine analysis was done by flow cytometry when cells were stimulated with SLTRiP protein (Fig 3C) T epitope containing peptides (Fig 3D). Our results showed an elevated number and percentage of monofunctional IFN-γ secreting T-cells in SLTRiP immunized mouse splenocytes (Fig 3D). An increase in the number and percentage of polyfunctional IFN-γ and TNF-α secreting cells or dual cytokine positive cells, corresponding to approximately 15% was also observed in SLTRiP immunized mice compared to control (Fig 3E and 3F). The results of percentage of cytokine positive cells was obtained from duplicate set, each set having four mice. Overall, we observed a shift towards higher polyfunctionality in cytokine-producing T-cells. These cytokines were secreted from same cells indicating the dual functionality of these cells. These cytokines aid in the elimination of parasites by enhancing phagocytic activity of macrophages, generating reactive oxygen intermediates (ROS) and Nitric Oxide (NO), and stimulating T-cell proliferation. The cytokine response produced in response to SLTRiP immnunization points at anti-parasitic properties of SLTRiP.

### 3.3 PbSLTRiP specific long lasting protective memory responses are induced in mice immunized with SLTRiP protein

For a peptide or protein to be an effective vaccine it should be able to generate durable memory responses. To analyze the generation and extent of memory induced in mice in response to SLTRiP immunization, mice were immunized following prime-boost regimen (Fig 4A). Three months after immunization the mice were challenged intravenously with 5000 infective wild type sporozoites each. Two days post challenge the mice was analyzed for parasite load in liver. The decrease corresponding to nearly 100 fold in 18srRNA copy numbers was observed in the liver of mice immunized with SLTRiP compared to control (Fig 4B). Simultaneously, in another group of immunized mice the blood parasitemia was monitored from day 3 after sporozoite challenge. A delay in pre-patent period of more than two days was observed in SLTRiP immunized mice compared to control (Fig 4C).

**IV).**
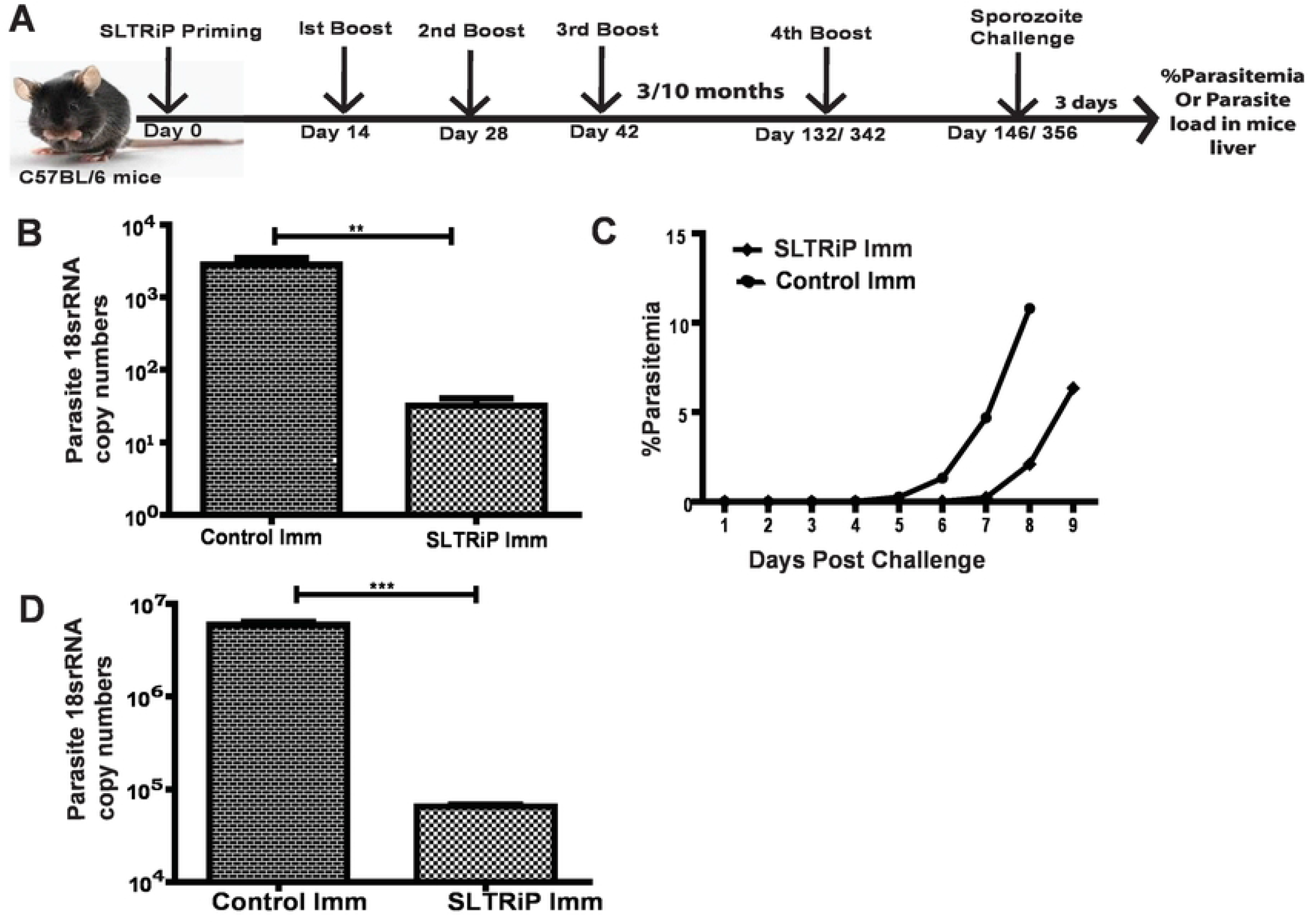
Long lasting Protective memory generation by SLTRiP immunization: Mice were immunized with SLTRiP, challenged with sporozoites and analyzed for liver stage parasite burden (A). The control was immunized with PBS emulsified in adjuvant. The exo-erythrocytic parasite burden was quantified from the liver of immunized mice 3 months (B). The Giemsa stained slides showing the pre-patent period of parasite in mice immunized with SLTRiP and control (C). The exo-erythrocytic parasite burden was quantified from the liver of immunized mice 10 months post immunization (D) All data are means and SEs based on six mice per group. **P < 0.01; ***P< 0.001 by Unpaired T-test. Control Imm: Control mice immunized with PBS: SLTRiP Imm: Mice Iimmunized with SLTRiP.

Another set of mice were immunized with SLTRiP following the prime-boost immunization regimen (Fig 4A); 10 months post immunization, mice were given another boost in the form of 5000 sporozoites under chloroquine treatment followed by challenge. Such an immunization mimics the occurrence of natural infection in endemic areas. The pattern of such immunization exposes the immune system of mice to native SLTRiP, which is a sporozoite and liver stage specific protein. We looked for the liver stage parasite burden of mice by quantitative PCR. The result shows more than 100 fold decrease in parasite 18srRNA numbers in the liver of SLTRiP immunized mice compared to control (Fig 4D). The delay in pre-patent period, and decreased parasite burden in liver in immunized mice, months after immunization indicates the generation of long lasting memory T-cells that are activated immediately after parasite exposure and are responsible for protection attributed to SLTRiP protein.

## Discussion

The liver stage of malaria is asymptomatic and cannot be detected until the parasite reaches blood stage. The liver stage represents a bottleneck during which the parasite multiplies manifold. A vaccine that targets parasite at this stage is highly desirable as it can reduce the disease morbidity. The ability of irradiated sporozoites to induce sterile protection mediated majorly by T-cells have been shown during 1960s [40]. The most successful subunit vaccine to date, RTS,S/AS01 has been shown to achieve up to 50% efficacy due the predominant induction of antibodies that block the sporozoite infection of hepatocytes. Alternatively, vaccination with many candidate subunit vaccines, viral vectors and DNA aim to induce CD8^+^ T cells capable of killing infected hepatocytes. The heterologous vaccination with many parasitic proteins also achieves protection that is correlated to the level of CD8^+^ T cells.

The kinetics of immune responses generated after vaccination, determine the functional attributes of a vaccine. Recent data highlight the importance of not only the quantity, but also the quality of the immune response induced [41]. The qualitative and quantitative mechanisms of effector T-cell responses, and the mechanisms by which they provide protection are poorly understood in case of *Plasmodium* infection. These maybe due to parasite’s phenotypic plasticity and absence of defined T-cell specific targets.

SLTRiP is a 45kDa molecular weight protein, which in its native form forms multimers. The amino acid sequence of protein shows the presence of low-complexity regions (LCRs). LCRs contain repeats of a single or short amino acid motifs, which are abundant in eukaryotes with unknown function. One hypothesis suggests that it may contribute to novel protein functions. SLTRiP has a tryptophan rich domain containing 19 conserved tryptophan residues, which likely interact with other proteins. SLTRiP is exported in host cell across the parasitophorous vacuole membrane where this tryptophan rich domain is likely involved in protein interactions, thus probably regulating many pathways of the host cell. The SLTRiP protein is of particular interest as it was able to elicit protective immune responses in mice model. Identification and characterization of epitopes on our target protein could help in the development of vaccines. In this context, the work aimed to determine the percentage of SLTRiP specific activated CD3^+^ T-cells generated in response to antigen stimulation. We observed that T-cells did recognize and respond to SLTRiP *in vitro* shown by the expression of activation marker on their surface. The quality of these cells in terms of protective efficacy has already been observed in SLTRiP immunization experiments [38, 39].

T-cells contribute to adaptive immunity by exerting pleiotropic effects that includes production of cytokines. These cells can release pro-inflammatory cytokines i.e IFN-γ, IL-2, and TNF-α, which are secreted following immunization and re-exposure to the pathogen antigens or in infectious disease state. Pro-inflammatory cytokines are associated with effective host defense against intracellular pathogens. These cytokines allow the accumulation of activated CD8+ T-cells and drive their differentiation into functional effectors. IFN-γ cytokine, especially is an important mediator for resistance against liver-stage of malaria. Infact IFN-γ mediated transient immunity against *Plasmodium* has been induced by immunization with radiation-attenuated sporozoites. We accessed the cytokine secretion and polyfunctionality of induced T-cell responses, as polyfunctional T-cells are correlated with protective efficacy.

The adaptive immune responses may cause generation of immunological memory due to which the immune system fights more rapidly and efficaciously to parasites encountered earlier. It also reflects the presence of antigen specific lymphocytes, which expand quickly on reexposure. Some of the efficient vaccines forming persisting memory cells are in use for many years now. These include vaccines against yellow fever, small pox, polio, DPT etc. The yellow fever vaccine, which is a live attenuated virus, gives lifelong protection. The memory T-cells have been observed even 35 years after vaccination [42, 43]. Memory B-cells generated against small-pox virus last >50 years in individuals [44]. Persistent memory B lymphocytes have been recognized for 5 or more years in vaccination with hepatitis B surface antigen against Hepatitis B Virus (HBV) [45]. This study also aimed to analyze protective efficacy of SLTRiP immunization months later. The decrease in liver stage parasite burden 10 months after immunization suggests the generation of durable and long lasting memory immune responses. Since the sporozoite boost under chloroquine was given only once, the memory recalled is attributed to SLTRiP recombinant protein. Ten months of mouse life, roughly corresponds to over 25 years of human life. The age correlation between human and mice, with an average lifespan of 60 years and 2 years respectively, suggests that one mice day is approximately equivalent to 30 human days. However, human and mice lifespan cannot be compared without considering different phases of the life. The precise correlation of the two must be drawn considering these phases separately. Nonetheless, determining the age relation in human versus mice helps in setting up experiments in murine models analogous to humans. Absence of a sterile protection giving vaccine against malaria is the major driving force behind identification of protective *Plasmodium* antigens. Although SLTRiP has certain limitations, like lack of sterile immunity and need for multiple doses, nonetheless, we show that SLTRiP elicited T cell responses that are important for protection in a pre-erythrocytic mouse model of malaria.

## Acknowledgement

We thank Mr. Vivek Kumar Pandey for his technical assistance in mosquito breeding, and Mr Kuldeep Chauhan for his help in analyzing FACS samples. This work received financial support from National Institute of Immunology (NII) core grant from Department of Biotechnology (DBT), Government of India, sanctioned to APS. A.Q. received financial support (JRF/SRF fellowship) from DBT through NII. This work was partly supported by a DST grant (EMR/2015/001546), Government of India sanctioned to APS.

## Data Sharing

The data that support the findings of this study are available from the corresponding author upon reasonable request.

## Abbreviations

*P.berghei*: *Plasmodium berghei*
PBS: phosphate buffered saline
RAS: Radiation Attenuated Sporozoites

